# Carbohydrate distribution via SWEET17 is critical for Arabidopsis inflorescence branching under drought

**DOI:** 10.1101/2023.01.10.523414

**Authors:** Marzieh Valifard, Azkia Khan, Rozenn Le Hir, Benjamin Pommerrenig, H. Ekkehard Neuhaus, Isabel Keller

## Abstract

Sugars Will Eventually be Exported Transporters (SWEETs) are the most recently discovered family of plant sugar transporters. Functioning as uniporters and thus facilitating the diffusion of sugars across cell membranes, SWEETs play an important role in various physiological processes such as abiotic stress adaptation. AtSWEET17, a vacuolar fructose facilitator, was shown to be involved in the modulation of the root system during drought. Moreover, overexpression of a homolog from apple results in increased drought tolerance of tomato plants. Therefore, SWEET17 appears to be essential for the plant’s drought response. Nevertheless, the role and function of SWEET17 in aboveground tissues under drought stress to date remains enigmatic. By combining gene expression analysis with analysis of the sugar profile of various aboveground tissues, we uncovered a putative role of SWEET17 in the carbohydrate supply, and thus cauline branch emergence and growth, particularly during periods of carbon limitation as occurs under drought stress. SWEET17 thereby being of critical importance for maintaining efficient reproduction under drought stress.

**Highlight:** The fructose transporter SWEET17 supports shoot branching by increasing mobilization of carbohydrates from vacuoles to supply the newly forming inflorescence branch, thereby maintaining efficient reproduction under drought stress.

## Introduction

As sessile organisms’ vascular plants are constantly exposed to a changing environment and such environmental conditions can alter either rapidly or gradually. Therefore, plants must constantly precept, react and adapt to their environment (Kleine et al., 2021; Schwenkert et al., 2022). As plant metabolism adapts, environmental conditions can affect important factors such as plant biomass accumulation and thus yield. One abiotic factor which markedly impairs plant growth and development is drought. Drought stress occurs for a variety of reasons, including low rainfall, high and low (below freezing) temperatures, high soil salinity or high light intensity. From an agricultural and physiological perspective, drought stress sets in when water availability decreases due to low soil moisture or when the rate of transpiration from leaves exceeds the water uptake by the roots (Salehi-Lisar and Bakhshayeshan-Agdam, 2016). Due to the projected global warming and climate change, the frequency and intensity of drought stress will increase worldwide (Dai, 2013; Basu et al., 2016; Bashir et al., 2021). Therefore, the probability of yield loss due to exceptional drought events will increase by about 20% in the future and already exceeds 70% for several crops such as soybean and corn (Leng and Hall, 2019). As diverse as the reasons of drought stress are the plants adaptive responses to this abiotic factor. E.g., plants react to water limitation with an array of morphological, physiological and biochemical adaptations, all following the general aim to maintain cell-homeostasis by decreasing cellular water depletion and/or increasing cellular water uptake. Such adaptations may include increased root formation, onset of stomata closure, relative decrease of shoot growth and sugar accumulation (Basu et al., 2016; Ahluweila et al., 2021; Bashir et al., 2021; Seleiman et al., 2021).

The accumulation of sugars has been well documented in a variety of metabolic and transcriptomic analyses under drought stress (Rizhsky et al., 2004; Cramer et al., 2007; Urano et al., 2009). It has been proposed that sugars then act as compatible solutes and decrease the water potential of the cell to maintain water retention and cell turgor (Krasensky and Jonak, 2012; Takahashi et al., 2020). In addition, sugars stabilize proteins and membranes (Hoekstra et al., 2001) and act as radical scavengers to maintain cellular redox balance under increasing accumulation of reactive oxygen species (ROS) promoted by drought stress (Miller et al., 2010; Kaur and Asthir, 2017). Accordingly, to fulfill their function as protein/membrane stabilizers and ROS quenchers, sugars need to be distributed throughout the whole plant system and in different subcellular compartments under abiotic stress (Pommerrenig et al., 2018; Keller et al., 2021). Thereby abiotic stresses, such as drought, lead to altered expression and activity of intra- and intercellular sugar transporters (Xu et al., 2018; Kaur et al., 2021).

Overall, the plant genome harbors numerous individual genes encoding carbohydrate-transport proteins that can be grouped in three major transporter families: the monosaccharide transporter-like (MST) family, the sucrose transporters (SUT/SUC), and the sugars will eventually be exported transporter (SWEET) proteins (Doidy et al., 2012; Pommerrenig et al., 2018; Wen et al., 2022). Of these three families, SWEETs are the most recently described transporter group (Chen et al., 2010) and to date, common features of all characterized SWEETs are their ability to mediate both influx and efflux of mono- and/or disaccharides at low sugar affinities (Chen et al., 2015a). SWEET transporters generally exhibit seven transmembrane domains and most SWEETs locate at the plasma membrane (Ji et al., 2022). However, three of them, namely SWEET2, SWEET16 and SWEET17 have previously been shown to localize to the tonoplast (Chardon et al., 2013; Klemens et al., 2013; Chen et al., 2015). Since the vacuole is the largest cellular organelle and because one of its main functions is the regulation of dynamic sugar storage and distribution, it does not surprise that especially vacuolar SWEET transporters show differential expression under abiotic stress conditions (Chardon et al., 2013; Klemens et al.,2013; Guo et al., 2014; Chen et al., 2015; Ji et al., 2022).

Recently, SWEET17, a vacuolar transporter with high specificity for fructose (Chardon et al. 2013; Guo et al., 2014), was shown to be involved in fructose-stimulated modulation of the root system under drought and thus directly involved in the plant’s drought response (Valifard et al., 2021). Since *SWEET17* expression is not only confined to the root region and high expression levels could also be found in above-ground tissue like the inflorescence stem (Guo et al., 2014), where its expression is explicitly confined to the vasculature (Chardon et al., 2013; Aubry et al., 2022), we focused on the role of the transporter in aboveground-tissues under drought stress. Therefore, we combined gene expression analysis with metabolite measurements of dissected Arabidopsis shoot tissues to reveal a possible involvement of SWEET17 in inflorescence branching under drought stress.

## Materials and Methods

### Plant cultivation and harvest

Wild types (Col-*0*) and two *sweet17* loss of function mutants (*sweet17-1* (SALK_012485.27.15.x) and *sweet17-2* (SAIL_535_H02); Chardon et al., 2013) were grown under different growth conditions based on the experimental design and purpose. For soil experiments, seeds were sown on standard soil (ED-73; Einheitserde Patzer; Sinntal-Altengronau, Germany) and plants were grown under short day conditions (10h light, 14h dark) with a light intensity of 125 μmol quanta m^-2^ s^-1^ at 21°C. To stimulate plants for initiation of the reproductive growth, four-week-old plants were transferred from short day to long day conditions (16h light, 8h dark) with the same light intensity and temperature as present at short days. For growth in hydroponic culture, seeds were germinated on germination medium, which was filled in detached lids from Eppendorf reaction tubes containing little holes, as described by Conn et al., (2013). The agar-filled lids were placed floating on plastic boxes containing liquid germination medium in a way, that developing roots can grow through the agar and extend directly to the liquid medium. After one week of growth, liquid germination medium was gradually exchanged with basal nutrient solution in the same composition as described in Conn et al., (2013). The basal nutrient solution was replaced weekly to ensure constant nutrient levels and pH of the medium. Plant material was harvested at the timepoints noted in the corresponding figure legends and if applicable was separated in the different aboveground-tissues leaf, stem, branch, flower and silique using a scalpel. Branch samples thereby represent first order lateral branches emerging from cauline leaf buds of the main inflorescence stem. For harvesting branch samples all inflorescences were removed from the branch. Plant material was directly frozen in liquid nitrogen after harvest and stored at −80°C until usage.

### Application of drought stress

To analyze the effects of drought stress on plant performance, drought was applied to the soil and hydroponic cultures based on different methods. For soil experiments, plants were exposed to drought conditions based on soil field capacity as explained in Valifard et al., (2021). Therefore, plants were kept at a determined water content in the soil, adjusted to a field capacity of either 100% (control) or 50%. The water content in the soil was checked and adjusted at 48h intervals until harvest. To apply drought stress in the hydroponic system, four-week-old plants were exposed to −0.5 MPa osmotic potential produced by polyethylene glycol 8000 (PEG 8000) according to Michel, (1983).

### Carbohydrate extraction and quantification

Frozen plant material was ground using a mortar and pestle. Carbohydrates were extracted as described in (Keller et al., 2021b). Briefly, 50 mg of pulverized plant material were extracted in 80% ethanol at 80°C for 30 minutes. After centrifugation (5min, 14000rpm), the supernatant was transferred into a new reaction tube and evaporated using a vacufuge concentrator (Eppendorf, Hamburg, Germany). The pellet remaining after evaporation was resolved in _dd_H_2_O. Sediments remaining from the carbohydrate extraction were washed with 80% ethanol and _dd_H_2_O twice and used for starch digestion. For that, 200 μl _dd_H_2_O were added to the washed pellet and samples were autoclaved for 40 minutes at 121°C. For hydrolytic cleavage of the starch, 200 μl of an enzyme mixture (5 U α-Amylase; 5 U Amyloglucosidase; 200 mM Sodium-Acetate; pH 4.8) were added to the autoclaved pellet and the mixture was incubated at 37°C for at least four hours followed by heat inactivation of enzymes at 95°C for ten minutes. Quantification of the extracted sugars (glucose, fructose, sucrose) and the hydrolyzed starch was performed using a coupled enzymic test (spectrophotometric analysis) as described in Stitt et al., (1989).

### Histological localization of *SWEET17*

The tissue localization of *SWEET17* was analyzed by histochemical analysis of transgenic plants, expressing the GUS (b-GLUCURONIDASE) reporter gene under control of the *SWEET17* promotor region (Valifard et al., 2021). Therefore, transgenic *ProSWEET17:GUS* plants were grown under short day conditions for four weeks and were transferred to long day conditions, followed by an application of drought stress at 50% FC for additional four weeks. Tissues of eight-week-old plants were stained by 5-bromo-4-chloro-3-indolyl-β-glucuronic acid (X-Gluc) solution according to Chardon et al., (2013) and the tissue localization of the *ProSWEET17:GUS* was documented using a Nikon SMZ1111 stereomicroscope combined with a ProgResC3 camera and the ProgResCapturePro 2.8 software (Jenoptik, Jena, Germany). To create thin sections of tissues, stained samples were dehydrated and embedded in Technovit 7100 resin (Kulzer, Hanau, Germany) as previously described by De Smet et al., (2004). Cross sections of four to 5.5 μm were prepared using a Reichert-Jung Biocut 2030 Microtome (Leica biosystems, Nußloch, Germany) and sections were observed as described above.

### Determination of reproductive growth parameters and yield

For determination of reproductive growth parameters and yield, total inflorescence height as well as the length of all first order cauline branches (emerging from cauline leaf buds of the main inflorescence stem) with a minimum length of 1 cm were measured on eight-week-old plants. For determination of the seed weight per plant, inflorescences of single plants were covered with paper bags as soon as all flowers turned to siliques. After ripening, seeds of single plants were harvested separately and the seed weight per plant was determined for ten individual plants of each line. From those ten individual plants, seeds of five plants were used to count and weight 500 seeds to determine the 500 seed weight.

### RNA extraction

Total mRNA was isolated from cauline branches and full rosettes of plants grown on soil and in hydroponic culture. Therefore, approximately 50 mg of ground tissue were extracted using the NucleoSpin RNA Plant Kit (Macherey-Nagel, Düren, Germany) following the user guidelines. Quality and quantity of the extracted RNA were photometrically checked using the NanoPhotometer N50 (Implen, München, Germany) and 1 μg of total mRNA was translated to cDNA using the iScript cDNA Synthesis Kit (Bio-Rad, Hercules, CA, USA) according to the instructions.

### Expression analysis via RT-qPCR

Analysis of gene expression was performed by quantitative real time PCR and was carried out in a CF X96™ real time cycler (Bio-Rad, Feldkirchen, Germany) using a standard two-step protocol with an annealing/elongation step at 58°C for 45 seconds. For quantification, the fluorescent dye iQ SYBR^®^ Green (Bio-Rad Laboratories, Feldkirchen, Germany) was used according to the manufacturer’s guidelines. The calculation of relative gene expression was performed using a modified 2^-ΔΔCT^ method (Livak and Schmittgen 2001). For transcript normalization the protein phosphatase 2A (PP2AA3; *AT1G13320*) and the SAND family protein (*AT2G28390*) were used as reference genes (Czechowski et al., 2005). Primers including their primer efficiencies used for expression calculation are documented as Supplementary Table 1.

### RNA Seq

RNASeq data was extracted from the dataset created for Khan et al. (2022, *preprint*). In brief, leaf discs from four-week-old Arabidopsis wild type plants were incubated in 3 mM MES containing 100 mM sugars (mannitol, glucose and fructose) overnight. The next day, leaf discs were harvested and stored at −80°C. RNA was isolated was using the NucleoSpin^®^ RNA Plant Kit (Macherey-Nagel, Düren, Germany) according to the manufacturer’s guidelines. RNA Sequencing then was performed and analyzed by Novogene (Cambridge, United Kingdom).

## Results

### *SWEET17* is transiently expressed in shoot tissue during drought stress

It is well known that plants accumulate sugars after drought exposure to mitigate the destructive effects of osmotic stress (Sami et al., 2016; Fulda et al., 2011). Therefore, especially tonoplast sugar transporters are stimulated under drought stress, leading to an increased sugar distribution via and an accumulation of sugars in the vacuole (Keller et al., 2021; Kaur et al., 2021). One of those vacuolar transporters is SWEET17, which’s expression was shown to be upregulated in specific root cells and involved in the initiation of lateral root development under drought stress (Valifard et al., 2021). While the role and function of SWEET17 under drought stress was recently described in the root tissue (Valifard et al., 2021), little is known about the function of the transporter in aboveground tissues under similar stress conditions.

We therefore aimed to elucidate the role of SWEET17 in the response of aboveground tissues to drought stress by conducting gene expression analysis under those stress conditions. To this end, wild types were grown in hydroponic culture for three weeks and treated with PEG 8000 to induce a controlled drought stress at an osmotic potential of −0.5 MPa and full rosettes were harvested at different time points during the treatment. It turned out that *SWEET17* transcript levels increased significantly in the shoot as early as one hour after the onset of drought (Figure 1 A), reached about three to four-fold abundance until twelve hours of stress, before returning to pre-stress levels after one day of drought treatment (Figure 1 A). Thus, *SWEET17* gene expression resembles the expression patterns of the known drought-induced vacuolar transporters *TST2* and *TST1* (Wormit et al., 2006) (Supplementary Figure S1).

**Figure 1:**
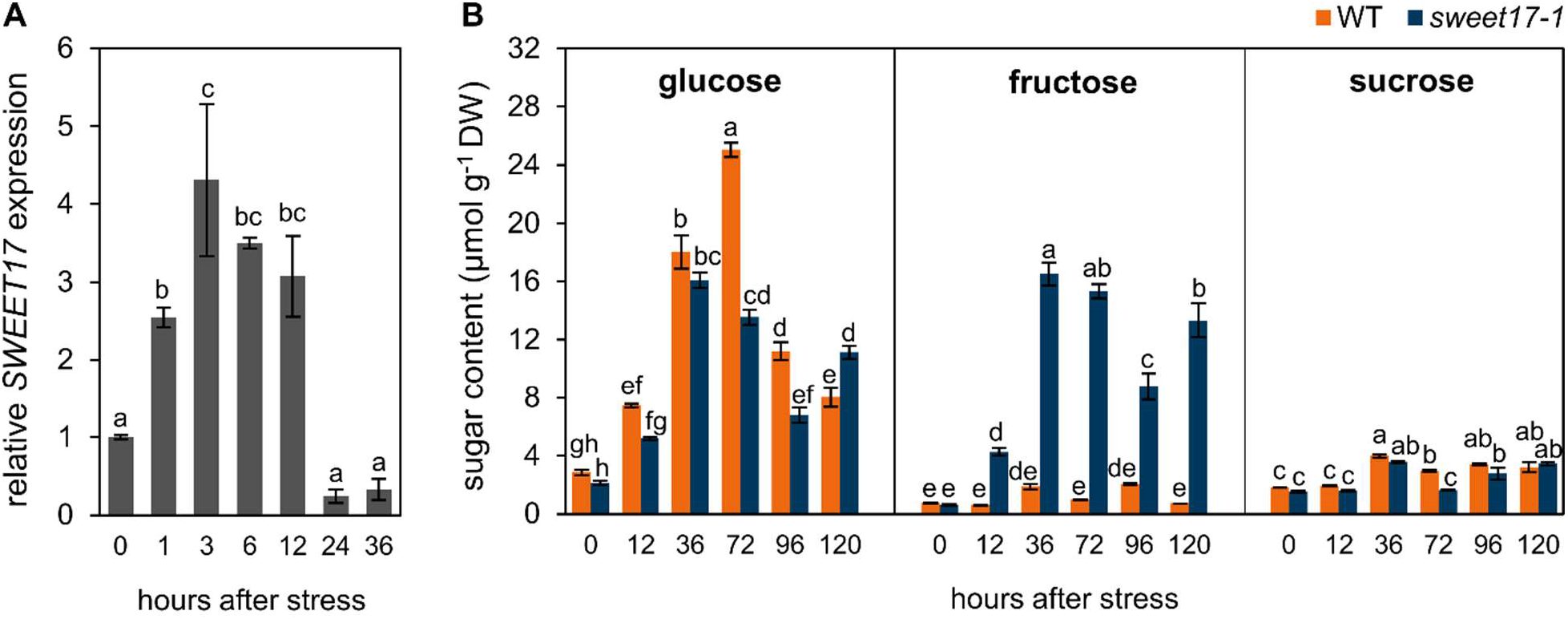
Drought-induced expression of *SWEET17* and response of *sweet17-1* mutant plants to short-term drought stress. Plants were grown in hydroponic system for three weeks and seedlings were exposed to artificial drought stress produced by PEG8000 (Ψs = −0.5 MPa). Full rosette tissue was harvested at different time points after onset of drought treatment and used for determination of gene expression and sugar quantification. A) Expression profile of *SWEET17* in wild type plants under short term drought stress. *SWEET17* expression was quantified in relation to *PP2AA3* and *SAND* expression and normalized on its expression in an unstressed control. Bars represent the mean from n = 3 biological replicates ±SE. Different letters indicate significant differences between timepoints according to one-way ANOVA with post-hoc Tukey testing (p < 0.05). B) Shoot sugar content of wild type and *sweet17-1* mutant plants grown under artificial drought stress. Bars represent the mean from n = 3 biological replicates ± SE. Different letters indicate significant differences between the different lines and timepoints according to two-way ANOVA with post-hoc Tukey testing (p < 0.05).

### *sweet17* mutants exhibit substantial accumulation of fructose during drought stress

Because SWEET17 activity is linked to fructose transport and sugar accumulation in roots under drought stress, we were interested to study corresponding effects in shoots. To investigate the effects of SWEET17 deficiency on the sugar content of aboveground tissues under drought, full rosettes of the well characterized *sweet17-1* knock out line (Chardon et al., 2013; Valifard et al., 2021) and wild type plants, grown in hydroponics under PEG 8000 induced drought stress, were analyzed (Figure 1 B).

Three-week-old rosette tissue of both, wild types and *sweet17* mutant plants, exhibited accumulation of glucose starting twelve hours after the addition of PEG. Glucose contents peaked at 36 and 72 hours after the onset of stress, followed by a decrease of glucose levels at 96 and 120 hours under drought in wild types and *sweet17-1* mutants, respectively (Figure 1 B). Although glucose content decreased after 96 hours of drought, it was still significantly higher than before onset of the stress treatment in both lines (Figure 1 B). Nevertheless, the accumulation of glucose in rosettes of wild types and *sweet17-1* mutants under drought stress was comparable (Figure 1 B). Unlike glucose, fructose accumulation could only be observed in rosettes of *sweet17-1* mutants under drought stress. At each analyzed time point after onset of drought, the fructose content was significantly higher in *sweet17-1* rosettes than in corresponding wild type samples (Figure 1 B). Starting from 0.76 μmol g^-1^ DW in wild types, fructose increased to a maximum of 1.87 μmol g^-1^ DW and 2.08 μmol g^-1^ DW after 36 and 96 hours under drought, respectively. In contrast, endogenous fructose content in the *sweet17-1* mutant was 0.63 μmol g^-1^ DW and increased to remarkable 16.55 μmol g^-1^ DW within the first 36 hours after onset of drought, a value nearly nine-times the wild type level (Figure 1 B). The levels of sucrose and starch showed comparable changes after exposure to drought stress as observed for glucose contents in wild types and the *sweet17-1* mutant. For both metabolites, a significant increase was observed in wild types and *sweet17-1* no later than 36 hours after onset of stress and remained high throughout the treatment (Figure 1 B and Supplementary Figure 2). Similar to glucose, sucrose and starch contents tended to be higher in wild types than in *sweet17-1* plants (Figure 1 B, Supplementary Figure S2). Nevertheless, differences in glucose, sucrose and starch contents between wild types and *sweet17-1* are minor compared to differences in the fructose content after drought stress (Figure 1 B).

### Loss of *SWEET17* results in altered sugar profiles in the inflorescence and branches

Since *SWEET17* expression was shown to be upregulated in aboveground tissues under drought stress (Figure 1 A) and since loss of *SWEET17* severely affects drought related sugar profiles (Figure 1 B), we wanted to investigate which of the aboveground tissues are most affected by *SWEET17* deficiency under drought stress. This analysis was of special importance since *SWEET17* expression is highest in the inflorescence stem (Guo et al., 2014). To this end, *sweet17-1, sweet17-2* and wild types were grown on soil under short day conditions for four weeks and subsequently transferred to long day conditions to initiate reproductive growth. With the shift in growth conditions, plants were subjected to 50% field capacity (FC) or 100% FC, respectively for four weeks afterwards. Eight-week-old plants then were subsequently dissected into the tissues: leaves, stems, branches, flowers and siliques prior to the extraction of sugars and starch (Figure 2). Our analysis revealed that overall glucose and sucrose concentrations were highest in the siliques of wild types, while glucose concentrations were lowest in the leaves and sucrose concentrations were lowest in branches (Figure 2 A and 2 C).

**Figure 2:**
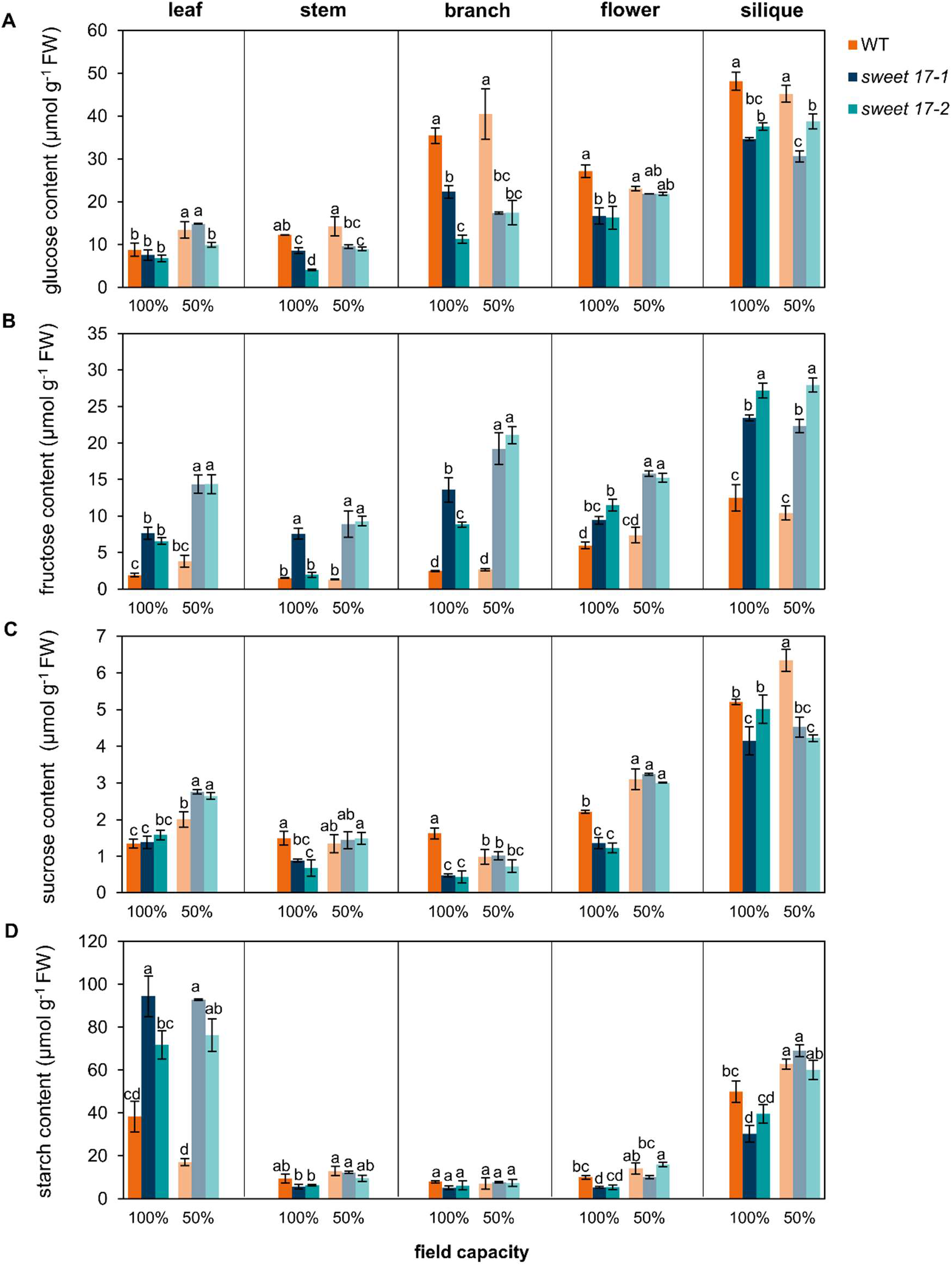
Sugar and starch content in aboveground tissues of Arabidopsis wild type and *sweet17* mutants in response to drought stress. Seeds were sown on soil and grown under short day conditions for four weeks and then were transferred to long day conditions followed by application of drought stress at 50% FC. After eight weeks of growth plants were dissected in leaf, stem, branch, flower and siliques and contents of glucose (A), fructose (B), sucrose (C) and starch (D) were measured. Starch contents were determined as hydrolyzed glucose. Bars represent the mean from n = 3 biological replicates ± SE. Different letters indicate significant differences between the different lines and conditions according to two-way ANOVA with post-hoc Tukey testing (p < 0.05).

As observed earlier, glucose contents were increasing in leaf-tissues upon drought treatment in wild types as well as in *sweet17* mutants and overall glucose level were comparable between all lines (Figure 2 A). In stems, branches, flowers and siliques no increase of glucose could be observed in any of the plant lines after growth at 50% FC. In addition, *sweet17* mutants exhibited lower glucose contents than wild types in those tissues (Figure 2 A). Regarding fructose, highest contents could always be observed in *sweet17* mutants, especially upon drought stress at 50% FC (Figure 2 B), while wild types did not show any accumulation of fructose under drought in any of the analyzed tissues (Figure 1 B, Figure 2 B). The biggest differences in fructose contents between wild type and *sweet17* plants could be observed in stems and branches under control, as well as under drought conditions.

Thereby, the concentration of fructose was already five- to 5.5-fold higher in *sweet17-1* stems and branches than in corresponding wild type tissues under unstressed conditions and increased to approximately seven-times the concentration of wild types under drought stress, indicating an important role of the transporter in these tissues particularly under drought (Figure 2 B).

Unlike fructose, sucrose contents were lower in stems, branches, flowers and siliques of *sweet17* mutants when compared to corresponding wild type tissues under unstressed conditions (Figure 2 C). In leaves, sucrose contents were comparable between the different plant lines (Figure 2 C). Drought stress led to an increase in sucrose contents in leaves and flowers of all tested lines, while in stems, branches and flowers sucrose concentrations increased solely in *sweet17* mutants, resulting in levels comparable to those of the wild types at 50% FC (Figure 2 C). In contrast to that, starch contents remained nearly unaffected by stress treatment. Only in siliques a significant increase in starch could be observed in all lines when exposed to drought (Figure 2 D). Overall starch levels were comparable between wild types and mutant plants except for the leaf tissue since mutant plants showed at least twice as high starch contents as present in wild types under each condition (Figure 2 D).

### *SWEET17* is expressed in the inflorescence stem during drought stress and branch formation

To investigate the tissue distribution of *SWEET17* in Arabidopsis, especially in stems and cauline branches of the inflorescence where differences in fructose contents between wild types and *sweet17* mutants are most pronounced (Figure 2 B), transgenic lines carrying the promotor region of the *SWEET17* gene fused to the β-glucuronidase reporter gene (*ProSWEET17:GUS*) were used (Valifard et al., 2021).

Histochemical localization of *ProSWEET17:GUS* in eight-week-old flowering plants demonstrated *SWEET17* promotor activity throughout the upper inflorescence under unstressed conditions and when plants were exposed to drought stress (50% FC; Figure 3 A and 3 B). Although the blue signal appeared throughout the whole upper inflorescence, *SWEET17* tissue localization analysis performed on cross sections of inflorescence stems revealed *SWEET17* promotor activity mostly in the xylary system along with faint signals in the cortex and the pith parenchyma (Figure 3 C). However, this distribution pattern was more pronounced in plants exposed to drought stress, with a strong blue signal observed in the pith region (Figure 3 D). Interestingly, when the cross-sections represented areas where branches connect to the main inflorescence stem, the blue GUS-signal could strongly be observed in the cortex in connecting areas of the outgrowing branch (Figure 3 E, as indicated by arrows). Latter staining pattern was also visible in plants exposed to drought stress (Figure 3 F), suggesting a role of SWEET17 in the formation of branches, especially where cortex parenchyma is re-differentiated to initiate meristematic cells required for branch development.

**Figure 3:**
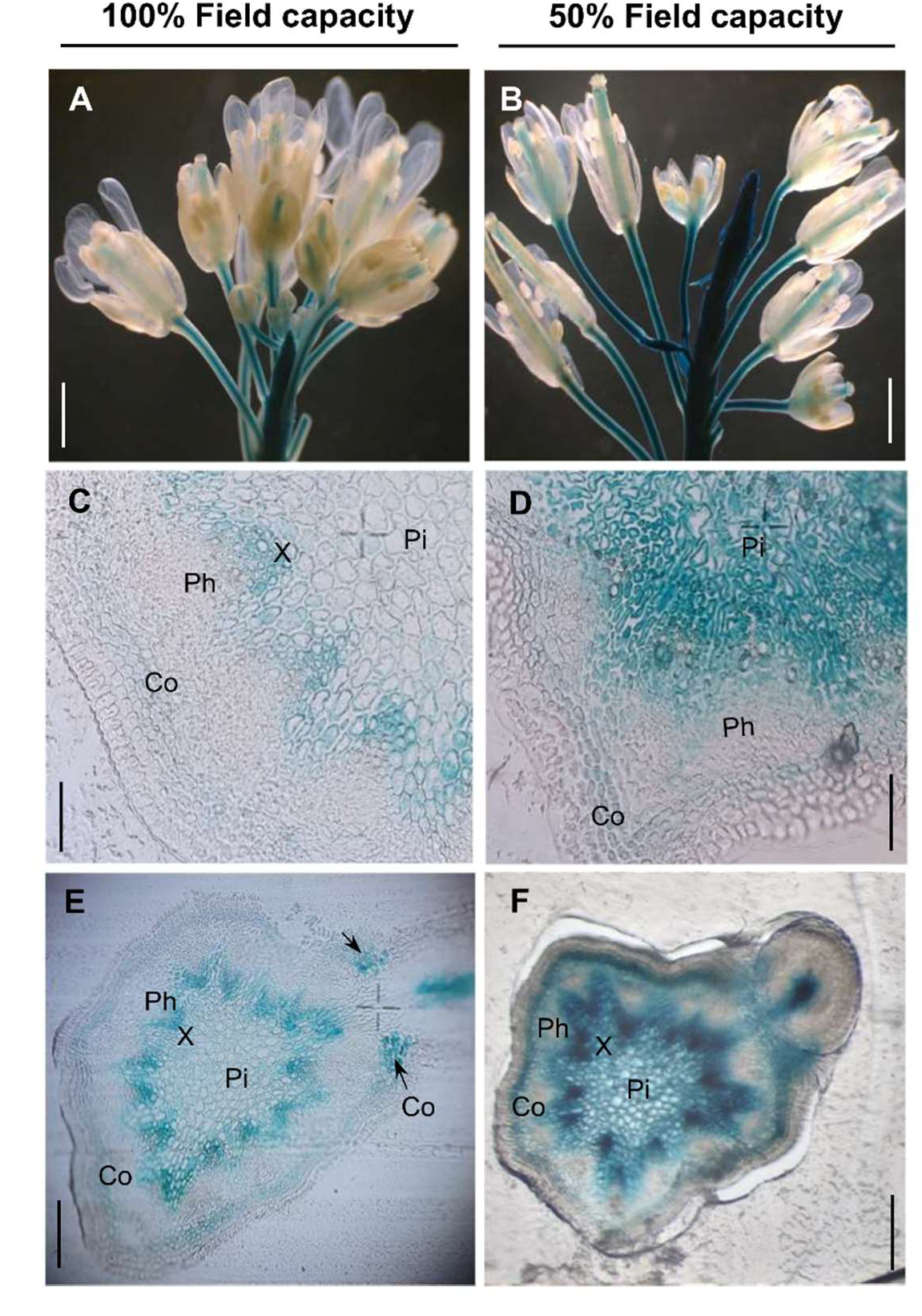
Expression pattern of *SWEET17* in Arabidopsis stem tissues. Histochemical localization of ProSWEETI7::b-GLUCURONIDASE (GUS) activity in eight-week-old inflorescence stem of Arabidopsis plants grown at 100% FC (A, C, E) and under drought stress at 50% FC (B, D, F). Seeds were sown on soil and grown under short day conditions for four weeks and transferred to long day conditions followed by application of drought stress at 50% FC. GUS signal was detected in the pith (Pi), xylem parenchyma (X) and cortex (Co) (C,D) especially in places of branch development (E, F) in eight-week-old plants. Pictures show representative staining from three individual plants grown at 100% and 50% FC, respectively. Inflorescence stems were embedded in resin (Kulzer Technovit 7100) and cross sections of 4μm (C, D, E) and 5.5μm (F) thickness were analyzed. Scale bars represent 250 μm (A, B), 30 μm (C, D), 100 μm (E, F).

### *sweet17* mutants exhibit decreased number of branches and lower seed yield per plant

Our analyses so far showed that *SWEET17* expression is drought induced and is present at sites of branch outgrowth (Figure 3). Interestingly, when comparing wild types and *sweet17* mutants under unstressed and especially drought stress conditions, cauline branches also showed most marked differences in their sugar composition (glucose and fructose) (Figure 2). Next, we analyzed effects of lacking SWEET17 activity on inflorescence morphology under control (100% FC) and drought conditions (50% FC).

We observed that mutant plants exhibited an overall shorter inflorescence under both, well-watered and drought stress conditions, when compared to corresponding wild type plants (Figure 4 A-C). However, under drought stress, wild type and mutant plants showed reduced inflorescence heights when compared to control conditions, with *sweet17* inflorescences being significantly shorter than that of wild types (Figure 4 C). Unlike inflorescence height, the number of branches did not differ between wild type and mutants under control conditions (Figure 4 D). When exposed to drought stress, the branch number was significantly reduced in all lines (Figure 4 D). However, this decrease was more pronounced in *sweet17* mutants (Figure 4 D). Similar behavior was observed for the branch length (Figure 4 E). While only *sweet17-2* mutants showed significantly shorter first order cauline branches than wild types at 100% FC, at 50% FC branch length was reduced in all lines. There, both *sweet17* mutant lines showed significantly reduced branch lengths when compared to the wild type (Figure 4 E). Therefore both, branch number and branch length appeared to be negatively affected in *sweet17* mutants under drought stress, as both parameters are comparable between all lines at 100% FC but are significantly lower in *sweet17* mutants under drought compared to corresponding wild types (Figure 4 D and Figure 4 E). Under control conditions, in *sweet17* mutants a lower inflorescence height resulted in a decreased seed yield per plant (Figure 4 F), while showing similar 500 seed weight (Figure 4 G) as wild types. A decrease of seed yield could also be observed in wild types under drought conditions (Figure 4 F). This negative effect was more severe in *sweet17* mutants (Figure 4 F).

**Figure 4:**
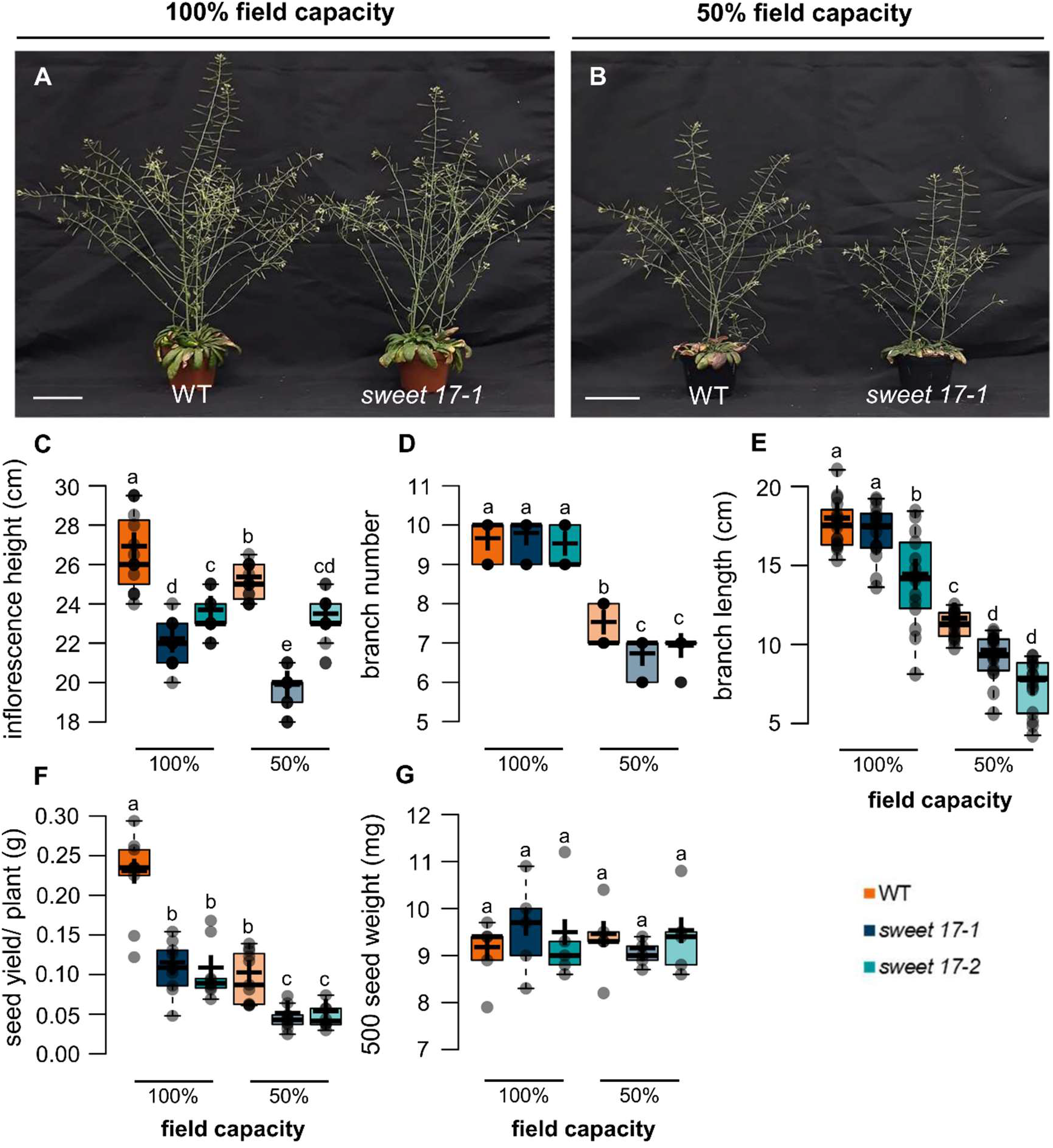
Phenotype of wild type and *sweet17* mutant lines exposed to drought stress at the reproductive stage. Comparison of inflorescence stem and branch development in *sweet17* mutant and wild type plants grown at 100% FC (A) and under drought conditions at 50% FC (B), as well as inflorescence height (C), branch number (D) and length (E), seed yield per plant (F) and 500 seed weight (G). Seeds were sown on soil and grown under short day conditions for four weeks and transferred to long day conditions followed by application of drought stress at 50% FC. Pictures were taken and reproductive parameters were analyzed on eight-week-old plants. Center lines in boxplots of reproductive parameters show the median, crosses represent the sample means. Box limits indicate the 25^th^ and 75^th^ percentiles and whiskers extend 1.5 times the interquartile range from the 25^th^ and 75^th^ percentiles. Datapoints of n = 15 biological replicates in (C-E), n=10 biological replicates (F) or n=5 biological replicates (G) are plotted as shaded circles. Different letters indicate significant differences between the different lines and conditions according to two-way ANOVA with post-hoc Tukey testing (p < 0.05). Scale bars represent 5cm (A, B).

### Expression of key regulators of branching and branch elongation are altered in *sweet17* mutants

The observation that *sweet17* mutants exhibit fewer and shorter branches, especially under drought stress (Figure 4), prompted us to investigate the expression of key transcription factors regulating both branch initiation and branch elongation (Figure 5). The process of branching is regulated by many different factors, including light phases, developmental stages, sugar availability and hormones like cytokinins and strigolactone (Rameau et al., 2015; Barbier et al., 2019). Accordingly, the complex regulation required for this process comprises different transcription factors.

**Figure 5:**
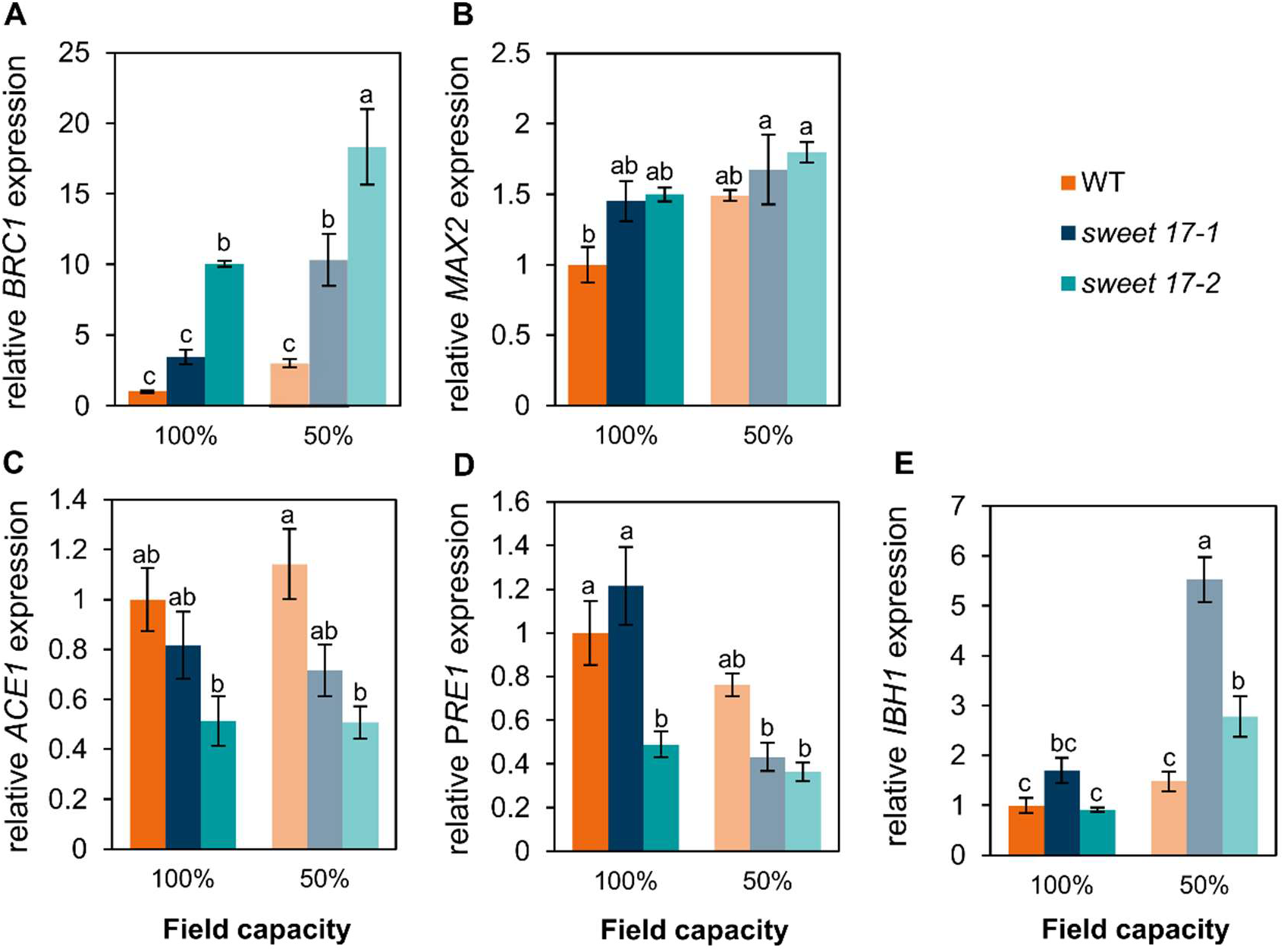
Expression profiles of central regulators of branching and branch elongation in wildtype and *sweet17* lines exposed to drought stress. For gene expression analysis seeds were sown on soil and grown under short day conditions for four weeks and transferred to long day conditions followed by application of drought stress at 50% FC. Lateral branches were cut from the main inflorescence stem of eight-week-old plants and used for RNA-extraction and following gene expression analysis. Expression of *BRC1* (A), *MAX2* (B), *ACE1* (C), *PRE1* (D) and *IBH1* (E) represents expression relative to *PP2AA3* and *SAND* and results were normalized on the expression of the WT under 100%FC. Values represent the mean of n=3 biological replicates ±SE. Different letters indicate significant differences between the different lines and conditions according to two-way ANOVA wit post-hoc Tukey testing (p<0.05).

One of these transcription factors is *BRANCHED1* (*BRC1*), an inhibitor of bud outgrowth that maintains bud dormancy (Aguilar-Martinez et al., 2019). Interestingly, *BRC1* expression was already higher in *sweet17-2* mutants compared with wild types under unstressed conditions and further increased under drought, resulting in significantly higher *BRC1* expression in *sweet17* mutants compared with the corresponding wild types (Figure 5 A). The expression of *BRC1* can be directly regulated via sugar availability or via sugar induced expression changes of upstream regulators like *MORE AXILLARY GROWTH2* (*MAX2*), which is involved in the strigolactone-dependent regulation of branching (Stirnberg et al., 2002). Drought in tendency induced the expression of *MAX2* in wild types (Figure 6 B). However, although not significant, *MAX2* expression was higher in *sweet17* than in wild type plants under both control and drought conditions (Figure 5 B).

**Figure 6:**
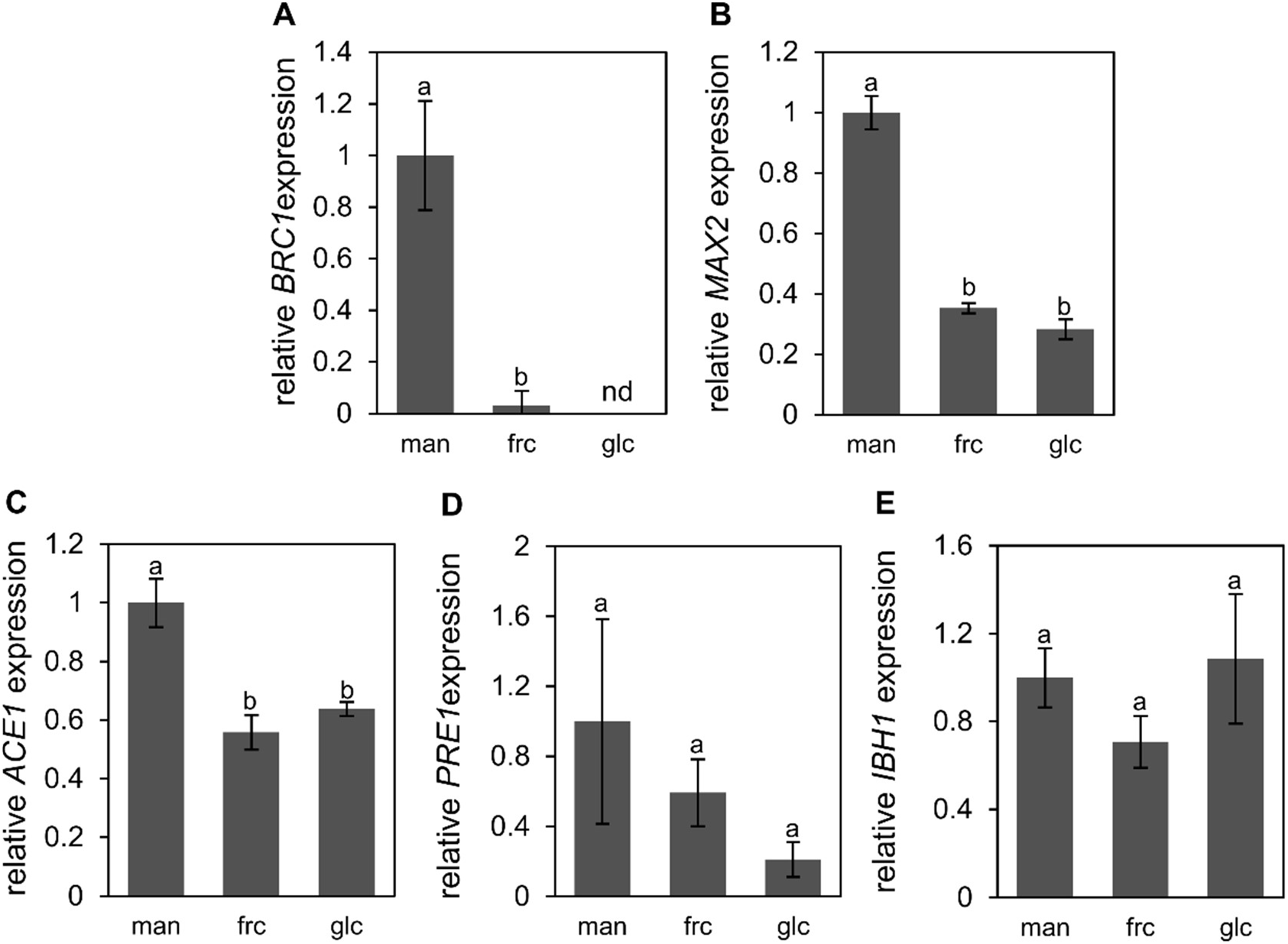
Expression of central regulators of branching and branch elongation in response to different sugars. Gene expression data was extracted from RNASeq performed by Khan et al. 2022. Therefore, leaf discs from four-week-old Arabidopsis plants were incubated in 3mM MES containing 100mM mannitol, 100mM fructose or 100mM glucose for 24 hours. Expression data in form of fpkm values of *BRC1* (A), *MAX2* (B), *ACE1* (C), *PRE1* (D) and *IBH1* (E) were extracted from the dataset and normalized on the fpkm value under mannitol treatment. Values represent the mean of n=3 biological replicates ±SE. Different letters indicate significant differences between the different conditions according to one-way ANOVA wit post-hoc Tukey testing (p<0.05).

In Arabidopsis, a series of bHLH (basic helix-loop-helix) transcription factors including ACTIVATOR FOR CELL ELONGATION1-3 (ACE1-3), PACLOBUTRAZOL-RESISTANT1 (PRE1) and INCREASED LEAF INCLINATION1 BINDING bHLH1 (IBH1) have been identified as regulators of cell elongation in response to environmental factors and developmental stages (Wang et al., 2018; Ikeda et al., 2012; Zhiponova et al., 2014). The expression of *ACE1* and *PRE1*, both being inducer of cell elongation, were already significantly reduced in *sweet17-2* mutants under control conditions (Figure 5 C and 5 D). Under drought, *ACE1* expression did not change in any of the analyzed lines, therefore *sweet17-2* still showing significantly reduced expression values when compared to the corresponding wild types (Figure 5 C). The expression of *PRE1* was significantly reduced in *sweet17-1* mutants under drought stress, while expression in wild types and *sweet17-2* was not affected. Therefore, although not significant, both mutant lines showed a lower expression of *PRE1* under drought stress when compared to wild types (Figure 5 C). Expression profiles of *IBH1* did not differ between *sweet17* mutants and wild types under unstressed conditions (Figure 5 E). However, under drought stress *IBH1* expression was significantly induced in both *sweet17* mutants leading to significantly higher expression values than in wild types (Figure 5 E).

### Expression of branching- and branch elongation regulators is influenced by monosaccharide levels

Given that the expression of key factors regulating branching and branch elongation differs between wild types and *sweet17* mutant plants, it is reasonable to speculate that fructose, the sole transport substrate of SWEET17 (Chardon et al., 2013), exerts a signaling function in the developmental process of branching. Therefore, to verify this hypothesis we extracted data from RNA-Seq analysis of Arabidopsis wild type leaf discs subjected to mannitol (as control) or fructose (Khan et al., 2022 *preprint*). In addition, extracted from the same dataset, we analyzed the effect of glucose, as another abundant monosaccharide, on global gene expression in Arabidopsis.

Expression of the three transcription factors *BRC1, MAX2* and *ACE1* were regulated by both fructose and glucose treatment, which generally led to a marked downregulation of their expression (Figure 6 A-C). Expression of *BRC1* was at such low levels after glucose feeding that it could not be detected in RNASeq analysis (Figure 6 A). Although not significant, *PRE1* showed an in tendency similar regulation with decreased expression levels after supply with either glucose or fructose (Figure 6 D). In contrast, *IBH1* expression was not affected by fructose and glucose treatment (Figure 6 E). Overall, a marked pattern of regulation of three of five key regulators of branching and branch elongation could be observed upon feeding external sugars.

## Discussion

The functions of sugars in plant metabolism are manifold. Sugars not only represent the main source for cellular energy and precursors of several important and abundant metabolites, but they also represent the major transport form of nutrients and energy, are involved in post-translational modification of proteins and lipids, play important roles in signal transduction, act as compatible solutes and represent efficient quenchers for reactive oxygen species (ROS). Latter abilities make sugars to important components of the complex plant stress resistance program (Ruan et al., 2014; Keller et al., 2021; Ji et al., 2022). Therefore, the ability to store and transport sugars intra- and intercellular is essential for the plants adaptation to its environment and has impact on the plants developmental processes (Wingenter et al., 2010; Klemens et al., 2013, Klemens et al., 2014; Patzke et al., 2019; Rodrigues et al., 2020).

Plant sugar sensing is a well described process and leads to the adjustment of the expression of a wide number of genes (Rolland et al., 2002, 2006). The best characterized plant sugar sensing system is represented by the sensor protein HEXOKINASE1 (Xiao et al., 2000; Moore et al., 2003), which connects changes of the cytosolic glucose concentration to altered transcription efficiency of nuclear located genes (Cho et al., 2006). Thus, it is not surprising that especially the subcellular composition of sugars affects the development of both, soil-located and aboveground plant organs (see e.g., Tjaden et al., 1994, Patzke et al., 2019, Valifard et al., 2021). Detailed analyses of various mutants revealed that especially the activity of vacuolar sugar transporters is important for the control of the cytosolic sugar levels (Wormit et al., 2006; Wingenter et al., 2010; Poschet et al., 2011; Klemens et al., 2013).

In line with these facts is the observation that the vacuolar fructose facilitator SWEET17 (Chardon et al., 2013; Guo et al., 2014) is critical for the initiation of lateral root formation, especially under drought stress (Valifard et al., 2021). SWEET17 was shown to be one of the 17 members of the SWEET-family in Arabidopsis and similar to SWEET2 and SWEET16, SWEET17 locates to the vacuolar membrane (Chardon et al., 2013; Klemens et al., 2013; Chen et al., 2015; Eom et al., 2015). Homologous genes of *AtSWEET17* were shown to be upregulated in other species in response to a range of environmental stress stimuli, including salt, osmotic and drought stress (Zhou et al., 2018; Lu et al., 2019). Therefore, it is not surprising that also *AtSWEET17* gene expression exhibits strong induction upon drought stress, regardless of the tissue analyzed (Figure 1 A; Valifard et al., 2021).

Since SWEET17 was shown to act as a fructose facilitator, loss-of-function of this transporter, which preferentially acts as a vacuolar exporter under unfavorable conditions (Guo et al., 2014; Chandran 2015), results in accumulation of fructose in the vacuole and thus a higher total cellular fructose content (Figure 1 B, Figure 2 B; Chardon et al., 2013). The opposite reaction was observed in SWEET17-overexpressor plants, which showed significantly lower fructose contents in comparison to the wild type, especially when grown in a challenging environment (Guo et al., 2014).

Similar to other stress stimuli, osmotic stress leads to homeostatic imbalances in plant cells quickly after its onset (Kollist et al., 2019). Consequently, plants must adapt to these challenging conditions, which occurs at various levels comprising alterations in morphology, metabolism and gene expression. To counteract the deleterious effects of severe drought stress, plants accumulate high levels of osmoprotective compounds, such as proline and various sugars to restore their osmotic balance (Gurrieri et al., 2020; Keller et al., 2021). This general response nicely fits with the observation that drought stressed Arabidopsis plants accumulate glucose and sucrose (Figure 1 B, Figure 2).

Interestingly, only *sweet17* mutants were able to accumulate fructose under drought stress conditions (Figure 1 B, Figure 2). As fructose accumulating in vacuoles mainly originates from sucrose cleavage via vacuolar invertases, those findings indicate an essential function of the vacuolar invertase during adaptation to drought. Latter conclusion is fully in line with the generally important function of vacuolar invertase for sucrose hydrolysis in Arabidopsis under various conditions (Vu et al., 2020). Moreover, as shown in mono- and dicot species, drought induces vacuolar invertase gene expression and the resulting enzyme activity is a critical element of the response to drought stress (Kakumanu et al., 2012; Chen et al., 2021). Vacuolar sucrose, cleaved by invertases, originates from an increased activity of vacuolar sugar transporters like TST1 and TST2, which’s expression is known to be upregulated under drought stress (Supplementary Figure 1; Wormit et al., 2006). In wild types invertase cleavage products glucose and fructose can sufficiently be exported from the vacuole via sugar porters like SWEET17 (Chardon et al., 2013; Valifard et al., 2021) and ESL1 (Yamada et al., 2010; Slawinski et al., 2021), while in the vacuole of *sweet17* plants fructose remains to be trapped to a higher extend, leading to the observed fructose levels (Figure 1 B, Figure 2 B).

Drought induced differences in fructose accumulation between wild types and *sweet17* plants as well as *SWEET17* expression are most pronounced in cauline branches (Figure 2 B, Figure 3 A and 3 B). These changes are in line with our and previous observations revealing high expression of *SWEET17* in the xylem parenchyma of the inflorescence stem (Figure 3 C; Guo et al., 2014). In those cells, SWEET17 is involved in the maintenance of fructose homeostasis to sustain the formation of xylem secondary cell wall (Aubry et al., 2022).

Both *SWEET17* transcript and SWEET17 protein were also detected in the cortex (Figure 3 C; Guo et al., 2014; Aubry et al., 2022; Hoffmann et al., 2022) and the pith (Figure 3 D; Hoffmann et al., 2022) of the stem, whereby expression of *SWEET17* in the pith is promoted under drought stress. Key characteristics of pith cells are their large size, large vacuoles and the fact that pith cells they are surrounded by vasculature (Lev-Yadun, 1994; Zhong et al., 2000), making them an ideal storage tissue. In addition, pith cells harbor a variety of sugar transporters, as e.g. carbohydrate transporters of the EARLY RESPONSE TO DEHYDRATION SIX-LIKE (ERDL) and SUCROSE TRANSPORTER (STP) families, known to be expressed in the inflorescence stem, show high expression levels in the pith of this organ (Shi et al., 2021; Dinant and Le Hir, 2022). Therefore, the pith and in particular its subcellular sugar distribution may play an important role in plant developmental processes, such as the development of the vascular system, a process that is clearly influenced by sugar signaling and thus by sugar availability and distribution (Dinant and Le Hir, 2022). In accordance with suggestions by Dinant and Le Hir (2022), increased *SWEET17* expression in the pith allows fructose to be mobilized from vacuoles and serve as carbohydrate source for local sinks such as the xylem tissue (Spicer 2014; Aubry et al., 2022). However, not only vascular tissue but also buds of cauline branches represent local sinks. Thus, increased expression of *SWEET17* could support bud outgrowth and branch development. This hypothesis gains support by strong *SWEET17* expression in the cortex, especially where branches are connected to the main stem (Figure 3 E and 3 F). This expression pattern resembles *SWEET17* expression in the outgrowing region of lateral roots where fructose specifically mobilized from vacuoles via SWEET17 is involved in controlled initiation of lateral root formation (Valifard et al., 2021).

During drought experiments *sweet17* mutants showed impaired biomass accumulation because of a limited water uptake ability due to lower root biomass (Valifard et al., 2021). Low water availability usually results in metabolic impairments such as decreased photosynthesis (Pinheiro and Chaves, 2011) and altered long distance transport (Keller et al., 2021). Together these factors result in inhibition of growth and therefore accumulation of sugars in leaf blades (Chaves and Oliveira, 2004) as observed in drought affected *sweet17* mutants (Figure 2). While sugars are synthesized and accumulate in source tissues, sink tissues like roots or siliques rely on carbohydrate distribution via the phloem, which is impaired under drought stress. In accordance, *sweet17* mutants accumulated higher carbohydrate contents in the leaves, while siliques showed lower contents of glucose and sucrose than the corresponding wild types under drought (Figure 2), highlighting a possible involvement of *sweet17* in carbohydrate distribution to branches and reproductive organs.

Shoot branching, like root development (Takahashi et al., 2013), is stimulated by the availability of sugars (Rabot et al., 2012; Mason et al., 2014; Barbier et al., 2015; Fichtner et al., 2017; Barbier et al., 2019). Sugar availability, as well as auxin-, strigolactone- and cytokinin levels exert influence on the process of branching (Thimann and Skoog 1933; Dun et al., 2012; Rameau et al., 2015; Balla et al., 2016; Dierck et al., 2016; Barbier et al., 2019). All these factors trigger changes in *BRANCHED1* (*BRC1*) expression, a central regulator of shoot branching, because it inhibits bud outgrowth and maintains bud dormancy (Aguilar-Martinez et al., 2019). *BRC1* expression under both, control and drought conditions were found to be higher in *sweet17* mutants when compared to corresponding wild types (Figure 5 A). This went along with a lower number of stem branches in mutant plants under drought (Figure 4 B and 4 D) and because *BRC1* expression was shown to be downregulated by external sucrose (Mason et al., 2014), it is reasonable to expect additional regulation by the supply of fructose and glucose, as shown above (Figure 6 A). In addition, the upstream integrator of the branching response named MORE AXILLARY GROWTH2 (MAX2) (Stirnberg et al., 2002), which’s expression is known to be negatively affected by sucrose (Barbier et al., 2015) and the external supply of glucose and fructose (Figure 6 B), was in tendency found to be increased in *sweet17* mutants (Figure 5 B). Thus, increased fructose mobilization by high expression of *SWEET17*, as occurring under drought, leads to suppression of *BRC1* and *MAX2*, which in sum stimulates branching. Because *MAX2* expression and signaling are directly influenced by factors such as strigolactone concentration (Chevalier et al., 2014; Khuvung et al., 2022) it is not surprising that observed *MAX2* expression differences between wild types and *sweet17* mutants are not significant (Figure 5 B). Anyhow *MAX2* expression supports the observed *BRC1* regulation and branching differences between wild types and *sweet17* mutants (Figure 4 and Figure 5 A).

Elongation and growth of plant cells is regulated by a tri-antagonistic series of helix-loop-helix (bHLH) transcription factors including ACTIVATOR FOR CELL ELONGATION1-3 (ACE1-3), PACLOBUTRAZOL-RESISTANT1 (PRE1) and INCREASED LEAF INCLINATION1 BINDING bHLH1 (IBH1) (Bai et al., 2012; Ikeda et al., 2012; Zhiponova et al., 2014; Wang et al., 2018). Expression of these factors differed significantly between wild types and *sweet17* mutants, especially under drought treatment (Figure 5 C-5 E). Thereby, reduced expression of the positive regulators *ACE1* and *PRE1* as well as an increased expression of the cell elongation inhibitor *IBH1* resulted in a decreased branch length in *sweet17* mutants, especially under drought stress (Figure 5 C-E). As the expression of the tri-antagonistic signaling cascade can be influenced by various environmental factors (Bai et al., 2012), it is reasonable to expect regulation of these genes by sugar availability, as for *ACE1* significantly and for *PRE1* in tendency observed above (Figure 6 C and 6 D). Sugar regulation of the corresponding signaling genes, as well as differences in their expression between wild types and *sweet17* plants reinforce the idea of a possible involvement of SWEET17 not only in shoot branching but also in branch elongation.

Overall, our results reveal high expression of *SWEET17* in the pith and cortex in areas of branch emergence of the inflorescence stem (Figure 1 A and Figure 3), high differences in the sugar profile of the main inflorescence stem and cauline branches between wild types and *sweet17* mutants (Figure 2), as well as differential expression of sugar regulated branching and branch elongation regulators (Figure 5 and Figure 6). In summary, these results suggest a supportive role of SWEET17 in shoot branching. Fewer branches as well as a limited branch length in *sweet17* mutants resulted in lower seed yield per plant, indicating an important function of SWEET17 for plant productivity. We believe that in wild types grown under drought conditions - in which sugar availability in sink tissues is limited by impaired photosynthesis and reduced functionality of the carbohydrate transport circuit (Li et al., 2017; Liang et al., 2020; Keller et al., 2021) - SWEET17 might lead to increased mobilization of sugars from the vacuoles of the pith to maintain carbohydrate supply to lateral bud formation. An idea supporting the assumption that SWEET proteins increase sugar mobilization to sink tissues during abiotic stress and therefore maintaining crop productivity (Anjali et al., 2021).

## Abbreviations

FC: field capacity
FW: fresh weight
DW: dry weight

## Supplementary Data

Fig. S1. Drought-induced expression of *TST2* and *TST1*.

Fig. S2. Drought-induced changes in starch content of *sweet17-1* mutant plants.

Table S1. List of Primers used in this study.

## Acknowledgements

We thank Ralf Pennther-Hager and Paul Berger for their help with plant cultivation, as well as Melissa Meinert, Valentyna Pliushch and Anastasiia Plakhotnia for technical assistance in the lab.

## Author Contribution

M.V., I.K. and A.K.: Investigation; H.E.N and M.V.: Conceptualization; M.V. and I.K.: Validation, Visualization; I.K.: Writing-original draft preparation; M.V., R.L.H., B.P. and H.E.N.: Writing-Review & Editing; H.E.N.: Supervision

## Conflict of interest

The authors declare no conflict of interest.

## Funding

This work was supported by a grant by the Alexander von Humboldt Foundation to M.V.. Financial support was also given to H.E.N. by the Federal State of Rhineland-Palatinate in the program “BioComp”.

## Data Availability

All data supporting the findings of this study are available within the paper and within its supplementary materials.

## Notes

### Competing Interest Statement

The authors have declared no competing interest.

