## Supplemental Data for "Carbohydrate distribution via SWEET17 is critical for Arabidopsis inflorescence branching under drought"

Valifard et al.

Fig. S1. Drought-induced expression of *TST2* and *TST1.*

Fig. S2. Drought-induced changes in starch content of *sweet17-1* mutant plants.

Table S1. List of Primers used in this study.

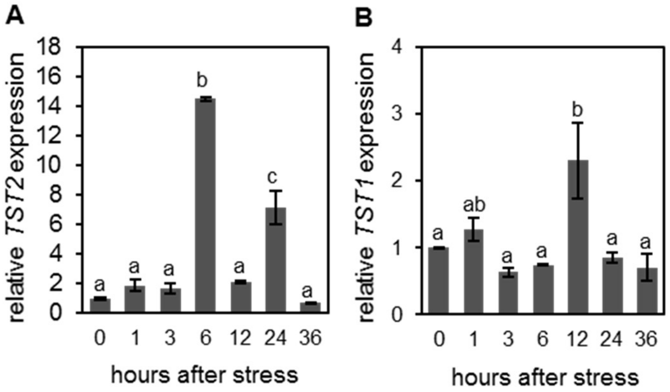


**Supplementary Figure 1: Drought-induced expression of *TST2* and *TST1* in wildtype plants.**

Plants were grown in hydroponic system for three weeks and seedlings were exposed to artificial drought stress produced by PEG8000 (Ψs = −0.5 MPa). Full rosette tissue was harvested at different time points after onset of drought treatment and used for determination of *TST2* (A) and *TST1* (B) expression. Gene expression was quantified in relation to *PP2AA3 and SAND* and normalized on the expression in an unstressed control. Bars represent the mean from n = 3 biological replicates ± SE. Different letters indicate significant differences between timepoints according to one-way ANOVA with post-hoc Tukey testing ($p < 0.05$).

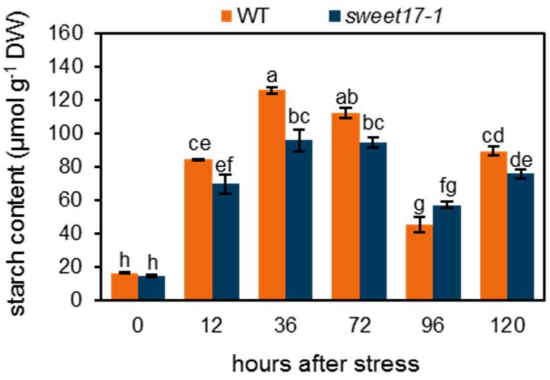


**Supplementary Figure 2: Drought-induced changes in starch content of *sweet17-1* mutant plants.**

Plants were grown in hydroponic system for three weeks and seedlings were exposed to artificial drought stress produced by PEG8000 (Ψs = −0.5 MPa). Full rosette tissue of WT and *sweet17-1* mutant plants was harvested at different time points after onset of drought treatment and used for starch quantification. Starch content was measured as µmol of hydrolyzed glucose per gram of dry weight. Bars represent the mean from n = 3 biological replicates ± SE. Different letters indicate significant differences between the different lines and timepoints according to two-way ANOVA with post-hoc Tukey testing ($p < 0.05$).

**Supplementary Table S1. List of Primer sequences and sequence information used in this study.**

| Primer | Acession Number | Sequence (5´-3´) | $R^2$ | Slope | Efficiency (%) |
|---|---|---|---|---|---|
| SAND family-fw | *AT2G28390* | AACTCTATGCAGCATTTGATCCACT | 0.998 | -3.2548 | 103 |
| SAND family-rev | | TGATTGCATATCTTTATCGCCATC | | | |
| PP2A subunit PDF2-fw | *AT1G13320* | TAACGTGGCCAAAATGATGC | 0.999 | -3.3607 | 98 |
| PP2A subunit PDF2-rev | | GTTCTCCACAACCGCTTGGT | | | |
| SWEET17-fw | *AT4G15920* | AGTGACAACAAAGAGCGTGAAATAC | 0.998 | -3.2419 | 103 |
| SWEET17-rev | | ACTTAAACCGTTGCTTAAACCACCC | | | |
| TST1-fw | *AT1G20840* | TTGCCGGCGAATTCTACTAAAGAG | 0.999 | -3.2011 | 105 |
| TST1-rev | | CAAGGATGGGACAATGCCACCA | | | |
| TST2-fw | *AT4G35300* | CATGGATCTTTCTGGTCGAAGGAC | 0.999 | -3.3893 | 97 |
| TST2-rev | | GATAAGACCGCGTGCACAATGC | | | |
| BRC1-fw | *AT3G18550* | AAAGCCAAGAAACCCAGCAG | 0.998 | -3.1275 | 109 |
| BRC1-rev | | GAGAAAGGGTTGTCGCGATC | | | |
| MAX2-fw | *AT2G42620* | CCATGGGGTCACACTCTTCT | 0.998 | -3.2983 | 101 |
| MAX2-rev | | GGAGCTTGATGTGGCGAATT | | | |
| ACE1-fw | *AT1G68920* | AATGGTGGATTCAAGCGACAGGAA | 0.998 | -3.2399 | 104 |
| ACE1-rev | | TGCTTGAGGGTCAGGTGTGA | | | |
| PRE1-fw | *AT5G39860* | CGTCGTTCTGATAAGGCATCAG | 0.999 | -3.3716 | 98 |
| PRE1-rev | | GACAGATTCGAGAAGCTGAGACA | | | |
| IBH1-fw | *AT2G43060* | GAAGGCTGCGTACGTTTCCA | 0.998 | -3.4107 | 96 |
| IBH1-rev | | CAAGAGGGCTCTGCTCCATA | | | |
